# Triple Arginines as Molecular Determinants for Pentameric Assembly of the Intracellular Domain of 5-HT_3A_ Receptors

**DOI:** 10.1101/561282

**Authors:** Akash Pandhare, Elham Pirayesh, Antonia G. Stuebler, Michaela Jansen

## Abstract

Serotonin type 3A receptors (5-HT_3A_Rs) are cation-conducting homo-pentameric ligand-gated ion channels (pLGICs) also known as the Cys-loop superfamily in eukaryotes. 5-HT_3_Rs are found in the peripheral and central nervous system, and they are targets for drugs used to treat anxiety, drug dependence, schizophrenia, as well as chemotherapy-induced and post-operative nausea and emesis. Decades of research of Cys-loop receptors have identified motifs in both the extracellular and transmembrane domains that mediate pentameric assembly. Those efforts have largely ignored the most diverse domain of these channels, the intracellular domain (ICD). Here we identify molecular determinants inside the ICD for pentameric assembly by first identifying the segments contributing to pentamerization using deletion constructs, and remarkably by making a small number of defined amino acid substitutions. Our work provides direct experimental evidence for the contribution of three arginines, previously implicated in governing the low conductance of 5-HT_3A_Rs, in structural features such as pentameric assembly.

## INTRODUCTION

Serotonin type 3A receptors (5-HT_3A_Rs) are cation-conducting homo-pentameric ligand-gated ion channels (pLGICs) also known as the Cys-loop superfamily in eukaryotes. 5-HT_3_Rs were first discovered in the peripheral nervous system in the gut (1). Only much later was it identified that agents developed for the treatment for chemotherapy induced emesis elicited behavioral effects in rodents indicative of a central nervous system effect. These receptors play a key role in the process of rapid excitatory neurotransmission in the human brain (2, 3). 5-HT_3A_Rs are targeted by many therapeutic drugs currently prescribed for the management of cancer chemotherapy-induced vomiting (4) as well as depression (5). They are also implicated as potential targets for the treatment of some neurological and psychiatric diseases and disorders (6). The Cys-loop superfamily additionally includes neuronal- and muscle-type nicotinic acetylcholine receptors (nAChRs), glycine receptors, and γ–aminobutyric acid type A (GABA_A_) receptors, which vary substantially in their ion-selectivity (7, 8). Cys-loop receptors assemble from five homologous subunits to form homo- or heteropentamers. Each subunit contains three separate structural and functional parts: 1. an N-terminal extracellular domain (ECD) mainly structured into two antiparallel β-sheets that harbors the ligand-binding site at subunit interfaces, 2. a transmembrane domain (TMD) consisting of four α-helical segments that shape the ion channel, and 3. a variable intracellular domain (ICD) (9). Recently, there has been remarkable progress in the determination of atomic resolution three-dimensional structures of a number of eukaryotic anion (10-13) and cation-conducting (14, 15) members of the Cys-loop superfamily. As a result, our structural understanding of subunit assembly, allosteric regulation, and conformational transitions associated with gating has improved significantly. Additionally, the structural basis underlying the actions of numerous clinically important drugs, which interact with the highly conserved ECD and/or TMD (11, 12, 16-18), and the complex interplay of conformational transitions between ECD and TMD associated with distinct functional state(s) of the channel have been elucidated (19-21).

While significant advances with regard to the structure and function of ECD and TMD have been achieved, structure determination of the ICD has remained elusive. The ICD of 5-HT_3A_Rs (5-HT_3A_-ICD) begins at the C-terminus of M3 as a loop (L1, Figure 1) and a short α-helix (MX) followed by a stretch of about 60 amino acids forming an intrinsically disordered region (L2) and a membrane-associated α-helix (MA). The MA helix forms a continuous helix with M4 at the cytoplasmic leaflet of the membrane bilayer. Pioneering electron microscopy (EM) studies on the cation-conducting *Torpedo* nAChR reported the presence of the MA helix in these receptors (22). Similar helical content pre-M4 is predicted for most other cation-conducting Cys-loop receptors but absent from anion-conducting superfamily members (Supplementary Figure 1). On the contrary, α-helical content corresponding to the MX helix is predicted for most anion- and cation-conducting channels. Overall, the lengths and sequences of ICDs are remarkably dissimilar between different subunits. The discovery that replacement of the ICD with a short linker for both anion- and cation-conducting Cys-loop receptors led to functional channels (23), inspired a series of studies utilizing the same or similar modifications for functional (24-28) and structural studies (14, 15, 29-33). Recent cryo-EM studies using full-length 5-HT_3A_Rs resolved L1-MX and the MA helix (20, 34), similar to a study where the ICD had been proteolyzed before crystallization (14). The inability to resolve L2 likely stems from its inherently ‘very dynamic’ nature (20). Therefore, structural – albeit incomplete - insights into the ICD are limited to *Torpedo* nAChR and 5-HT_3A_ receptors, and entirely lacking for anion-conducting Cys-loop receptors. Even though full-length GABA_A_ constructs were used for recent cryo-EM structure determination only a few amino acids of the ICD without defined secondary structure could be resolved (10, 13, 21). This provides the first experimental indication for the ICD of anion-conducting pLGICs being significantly different from cation-conducting channels.

**Figure 1.**
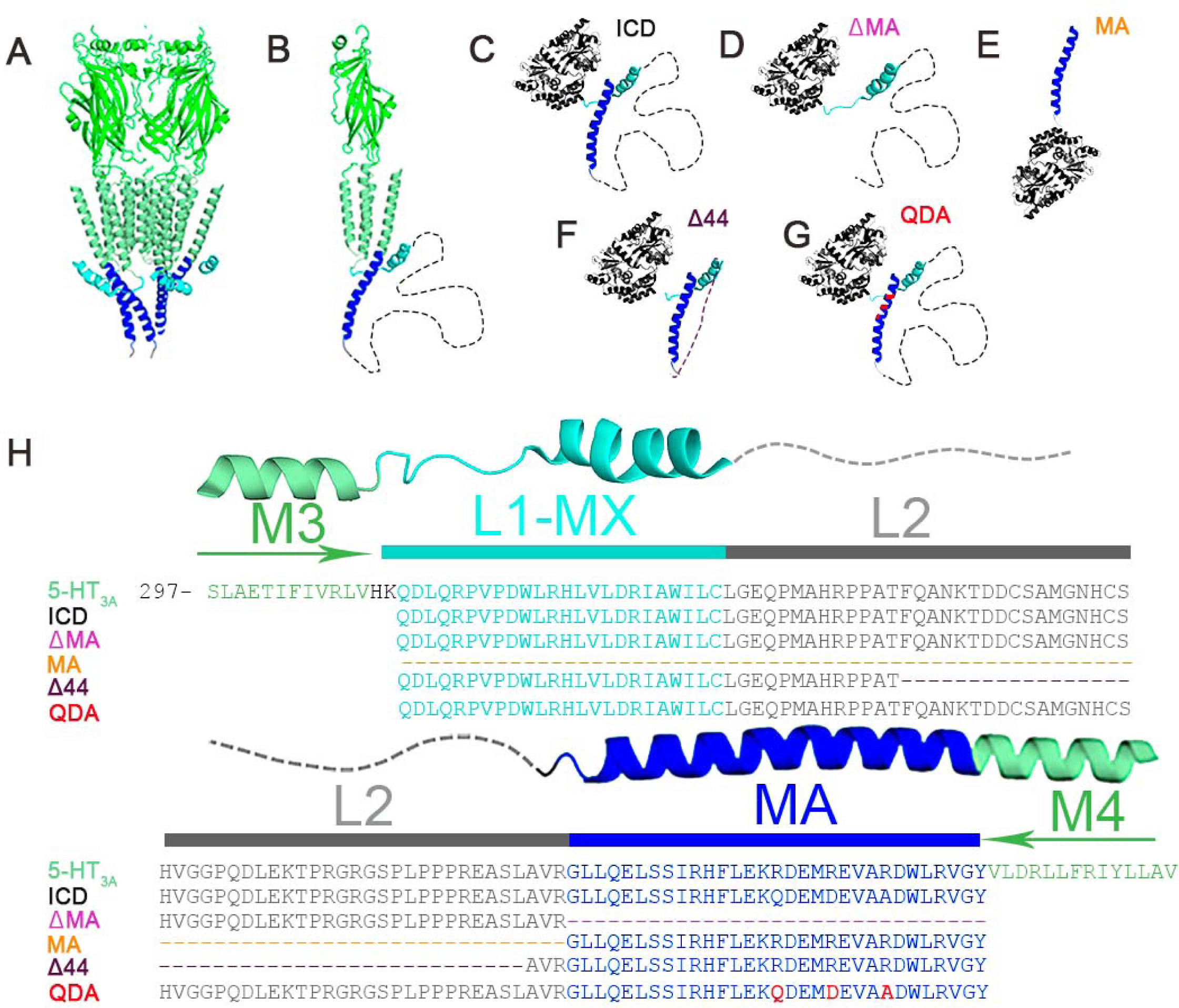
Structural representation and alignments of chimeras. **A)** Cartoon representation of the pentameric m5-HT_3A_ receptor X-ray structure viewed parallel to the plane of the membrane. ECD and TMD in green, MA and MX-helices in blue and cyan. **B)** A single subunit of the 5-HT_3A_ receptor viewed parallel to the membrane. The 62 residues not resolved in L2 are depicted as a dashed line. **C)** Cartoon representation of MBP-5HT_3A_-ICD-WT. The cytoplasmic domain of 5-HT_3A_R is fused to the C-terminus of MBP and serves as the template for the chimeras. **D)** The chimera MBP-ΔMA has the entire MA-helix removed. **E)** MBP-MA has the MA-helix attached to the C-terminus of MBP. **F)** MBP-Δ44 contains a 44 amino acid deletion in L2. **G)** MBP-QDA contains a triple mutation in its MA-helix and no deletions. **H)** Multiple sequence alignment of 5-HT_3A_R and the ICD chimeras highlighting deletions (dashed lines) or mutations (red).

The ICD is involved in determining ion conductance (23, 35) and modulation by drugs (36). Additionally, it interacts with various entities of the intracellular apparatus, which facilitate receptor sorting, assembly, trafficking, and anchorage (37, 38). Studies of subunit assembly and oligomerization have investigated the individual contributions of ECD (39, 40) as well as TMD (30, 31) at the molecular level. While both ECD and TMD can assemble individually into pentamers (29, 31, 41, 42), formation of hexamers of the ECD alone for GLIC (43) and non-native stoichiometries for α2β4 nAChR (31) exemplify that additional drivers outside these domains may mediate precise subunit stoichiometries and arrangements observed in nature. Intriguingly, this points toward the highly-diverse ICD as a mediator for defined oligomerization and also pentamerization in general. Indeed, we showed for the first time that the 5-HT_3A_-ICD alone can assemble into highly stable pentamers in the absence of the ECD and TMD (44). This remarkable experimental observation led us to investigate and uncover the molecular determinants for pentameric assembly of the 5-HT_3A_-ICD.

In the present study, based on a construct of the 5-HT_3A_-ICD in the absence of ECD and TMD, we introduced defined deletions over the entire span of the ICD, and finally a set of amino acid substitutions to determine the motif necessary for its pentameric assembly.

## MATERIALS AND METHODS

### Materials

BL21-CodonPlus-(DE3)-RIPL cells (Agilent Technologies, Santa Clara, CA); ampicillin (Fisher Scientific, Fair Lawn, NJ); chloramphenicol (Fisher Scientific, Fair Lawn, NJ); IPTG (Fisher Scientific, Fair Lawn, NJ); leupeptin (AdipoGen Life Sciences, San Diego, CA); pepstatin (AdipoGen Life Sciences, San Diego, CA); PMSF (Research Products International, Mt. Prospect, IL); TCEP-HCl (Oakwood Chemical, N. Estill, SC); lysozyme (MP Biomedicals, Solon, OH); Protease inhibitor cocktail III (Research Products International, Mt. Prospect, IL); DNAse I (Alfa Aesar, Ward Hill, MA).

### Molecular biology

The intracellular domain of the mouse 5-HT_3A_ receptor (accession number: Q8K1F4) was generated as a fusion construct with N-terminal maltose binding protein (MBP) as we described previously (45) using the pMAL-c2x (New England Biolabs) variant pMALX (46). Based on this MBP-5-HT_3A_-ICD template containing the entire wild-type ICD, deletions and substitutions were generated using the QuikChange II Site-Directed Mutagenesis kit (Agilent Technologies), and confirmed by DNA sequencing (GENEWIZ, South Plainfield, NJ). The resulting amino acid sequences for all constructs are displayed in Figure 1. We will refer to the construct with the full-length ICD as ICD. The construct with the MA-helix deleted will be referred to as ΔMA, the construct only consisting of the MA-helix will be MA, the construct with 44 amino acids between MX and MA removed will be Δ44, and finally the construct with three arginines mutated will be referred to as QDA.

### Expression and purification of the MBP-5-HT_3A_-ICD constructs

All plasmids for expression of MBP-5-HT_3A_-ICD fusion constructs with wild-type or engineered ICD were transformed into *Escherichia coli* (*E. coli*) BL21-CodonPlus-(DE3)-RIPL cells (Agilent Technologies). Cells were grown in Terrific Broth (TB) medium (1.2% (w/v) tryptone, 2.4% (w/v) yeast extract, and 0.4% (v/v) glycerol) supplemented with ampicillin (100 µg/ml), chloramphenicol (34 µg/ml) and 0.2% (w/v) sterile glucose, at 37°C and 250 rpm, in a shaking incubator. All concentrations indicated are final concentrations unless otherwise stated. At OD_600_ of 0.4-0.5, the cultures were induced by adding 0.4 mM isopropyl β-D-thiogalactoside (IPTG) and allowed to continue growth at 18°C for an additional 8 h. The cells were harvested by centrifugation at 4,600 *g* and 4 °C for 15 min, and then resuspended in (10 ml buffer/gram of the cell pellet) buffer A (20 mM Tris, pH 7.4, 200 mM NaCl, 1 mM TCEP, 2 mM EDTA) enriched with a freshly prepared protease inhibitor cocktail containing leupeptin (10 µg/ml), pepstatin (10 µg/ml), and 1 mM PMSF. After the cells were disrupted by treatment with lysozyme (100 µg/ml) and a freeze-thaw cycle, the lysate was clarified by ultracentrifugation at 100,000 *g* and 4°C for 1 h. The supernatant was immediately passed through a 0.2 µm pore-size bottle-top filter and loaded onto a pre-equilibrated amylose-resin column. Unbound proteins were washed off the column using 30 bed-volumes of buffer A, followed by elution with 20 mM maltose. Fractions were analyzed after separation in stain-free precast SDS-PAGE gels (4-20% Mini-PROTEIN^®^ TGX Stain-Free™, Biorad). These stain-free gels contain trihalo compounds that undergo UV-induced covalent modification of tryptophan residues for subsequent fluorescent detection. Fractions containing purified protein were pooled, and concentrated (up to approximately 5 mg/ml) using a 50 kDa molecular weight cut-off (MWCO) or a 100 kDa MWCO centrifugal filter (Amicon^®^ Ultra-15, Merck Millipore Ltd., Tullagreen, Ireland) for an additional purification step by size-exclusion chromatography (SEC).

### Size exclusion chromatography (SEC)

The amylose column purified, and concentrated protein samples were passed through an ENrich™ SEC 650 10 × 300 High-Resolution column (Biorad) pre-equilibrated with buffer B (20 mM HEPES, 150 mM NaCl, 1 mM TCEP, 5 mM Maltose, 0.01% NaN_3_, pH 7.4) for additional purification as well as molecular mass determination. For the apparent molecular mass estimation of each 5-HT_3A_-ICD construct (wild-type, deletion and substitution constructs), the SEC column was calibrated using thyroglobulin 669 kDa, ferritin 440 kDa, aldolase 158 kDa, conalbumin 75 kDa and ovalbumin 44 kDa according to the instruction manual (GE Healthcare). The column void volume (*V*_o_) was established with Blue Dextran 2000. The molecular mass of each construct was determined based on the calibration curve of the gel-phase distribution coefficient (*K*_av_) versus log molecular weight (log *M*_r_). The gel-phase distribution coefficient is calculated by the equation *K*_av_ = (*V*_e_-*V*_o_) / (*V*_c_-*V*_o_), where *V*_e_ = elution volume, *V*_c_ = geometric column volume and *V*_o_ = column void volume. The purified protein samples obtained after the final SEC step were analyzed by stain-free precast SDS-PAGE gels (4-20% Mini-PROTEIN^®^ TGX Stain-Free™, Biorad).

### SEC coupled with multi-angle light scattering (SEC-MALS)

We configured a Biorad BioLogic DuoFlow™ 10 system (including an F-10 workstation, auto sample inject valve, column holder and a UV detector with 254/280 nm filters) coupled with a miniDAWN-TREOS static 3-angle laser light scattering detector, and an Optilab-rEX refractive index (RI) detector (Wyatt Technology, Santa Barbara, CA) for conducting the SEC-MALS experiments. 250 μl of purified protein sample (0.1 – 0.5 mg) was passed, at a constant flow rate of 0.5 ml/min, through an ENrich™ SEC 650 10 × 300 High-Resolution column (Biorad) thoroughly pre-equilibrated with buffer B at room temperature. UV absorbance was measured with the detector at 280 nm. Light scattering and refractive index were monitored at a wavelength of 658 nm. BioLogic DuoFlow™ software version 5.3 (Biorad) was used to control the chromatography system, and the Astra 5.3.4 software (Wyatt Technology) was used for data collection and analysis. The light scattering detectors were normalized with monomeric bovine serum albumin (BSA; Sigma). Baseline settings for all laser detectors as well as peak alignment and band broadening correction between the UV, LS, and RI detectors were performed using Astra software algorithms. Data was processed to determine the weight-average molar mass and polydispersity of the protein sample using the Debye model as per the manufacturer’s instructions. Samples consisting of MBP alone were analyzed by SEC-MALS indicating that MBP was monomeric in solution (data not shown), consistent with previous small-angle X-ray scattering study (47). A minimum of three repeat runs were conducted for each 5-HT_3A_-ICD construct under identical experimental conditions.

## RESULTS

### Overview of approach

We earlier expressed and purified the intracellular domain of 5-HT_3A_Rs in fusion with a maltose-binding protein (MBP-5-HT_3A_-ICD; wild-type chimera), construct ICD, and showed that the domain self-associates, non-stochastically, as a pentamer in solution (44, 45). To gain further insights into the molecular mechanisms and interactions that govern pentameric assembly of the 5-HT_3A_-ICD, we adopted a systematic approach wherein a series of MBP-5-HT_3A_-ICD fusion constructs, bearing either large deletions or a set of amino acid substitutions within the ICD, were engineered. Our nomenclature for the different constructs is illustrated in detail in Figure 1. In brief, five constructs were investigated in this study: the initial full-length ICD containing 5-HT_3A_-ICD construct consisting of all 115 amino acids of the ICD; three deletion constructs with one lacking 44 amino acids from the disordered flexible region between the MX and MA helices (Δ44), one lacking the MA helix (ΔMA) and one consisting of just the MA helix alone (MA); and one construct based on the full-length ICD with three amino acids in the MA-helix mutated (QDA). All constructs were expressed in *E. coli* and purified to homogeneity. We then utilized a combination of biophysical and biochemical methods, including denaturing gel electrophoresis, size exclusion chromatography (SEC), and SEC in-line with multi-angle light scattering (SEC-MALS) to investigate the oligomeric state of each of the chimeras.

### Expression and purification of wild-type and engineered 5-HT_3A_-ICD constructs

Cytosolic expression in *E. coli* cells of the 5-HT_3A_-ICD as a chimeric protein containing an N-terminal MBP was under the control of a P_tac_ promoter (48). Robust soluble expression of the protein was achieved by carefully optimizing the conditions for culture induction and incubation as thoroughly described elsewhere (45). Subsequently, approximately 20 – 30 mg of ICD could be purified per liter of *E. coli* cell culture. Modifications (point or deletion) of this original construct, engineered using a combination of QuikChange and PCR methods (Figure 1), were also produced in a similar fashion with yields of these proteins being 8 – 30 mg/L of *E. coli* culture.

Tandem purification was accomplished on an Amylose Resin affinity column followed by size exclusion chromatography (SEC). SDS-polyacrylamide gel electrophoresis (SDS-PAGE) indicated a typical purity of the proteins of > 90% and > 95% after Amylose affinity chromatography and SEC, respectively (Figures 2 and 3).

**Figure 2.**
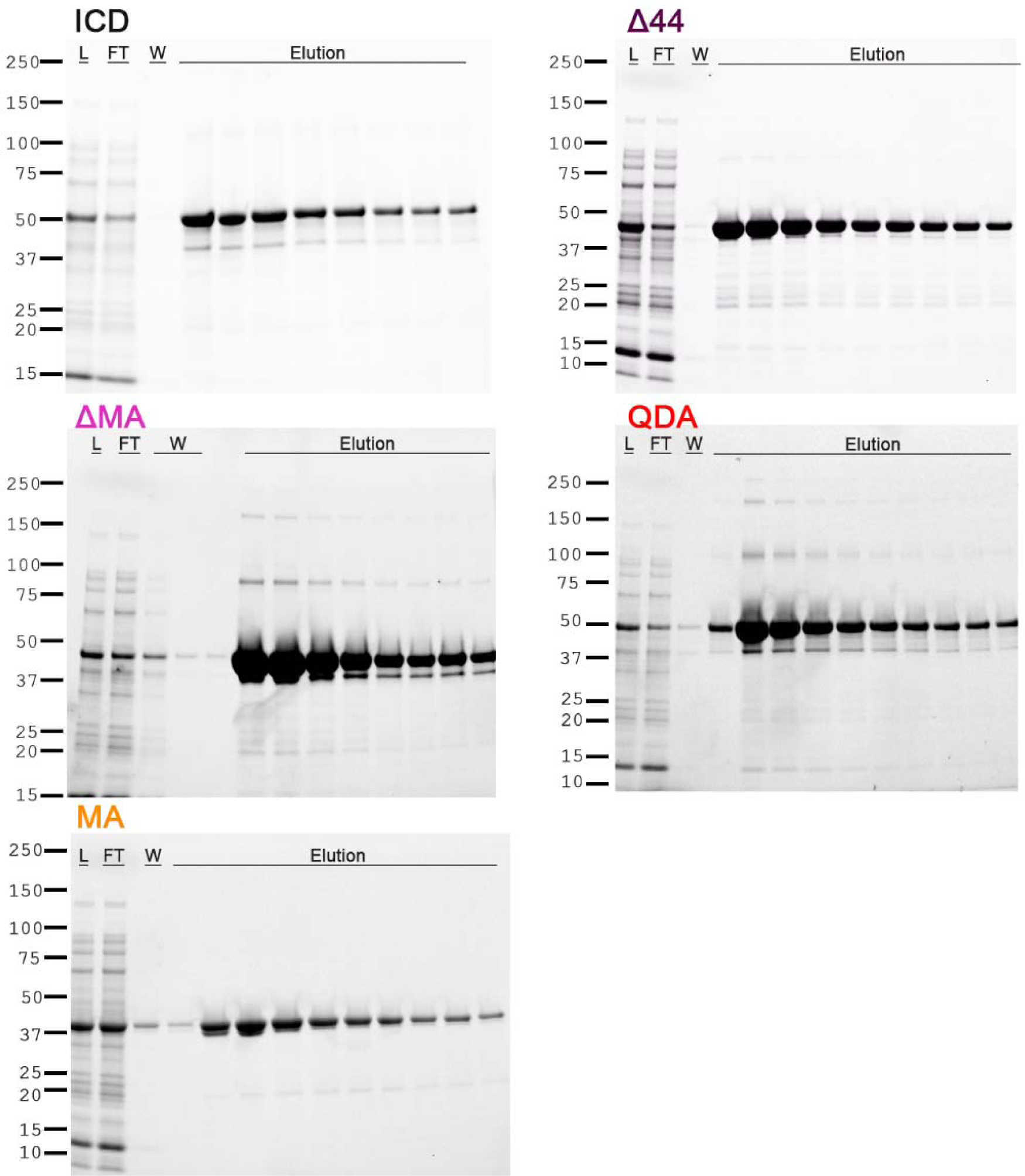
Affinity column purification of the ICD chimeras. The chimeras were purified by amylose-resin column chromatography. The soluble protein was loaded onto columns (L-loading material; FT-flow-through), followed by washes (W), and eluted. SDS-PAGE (4-20% Mini-PROTEIN® TGX Stain-Free™, Biorad) indicates the quality of purified protein and identifies peak fractions.

**Figure 3.**
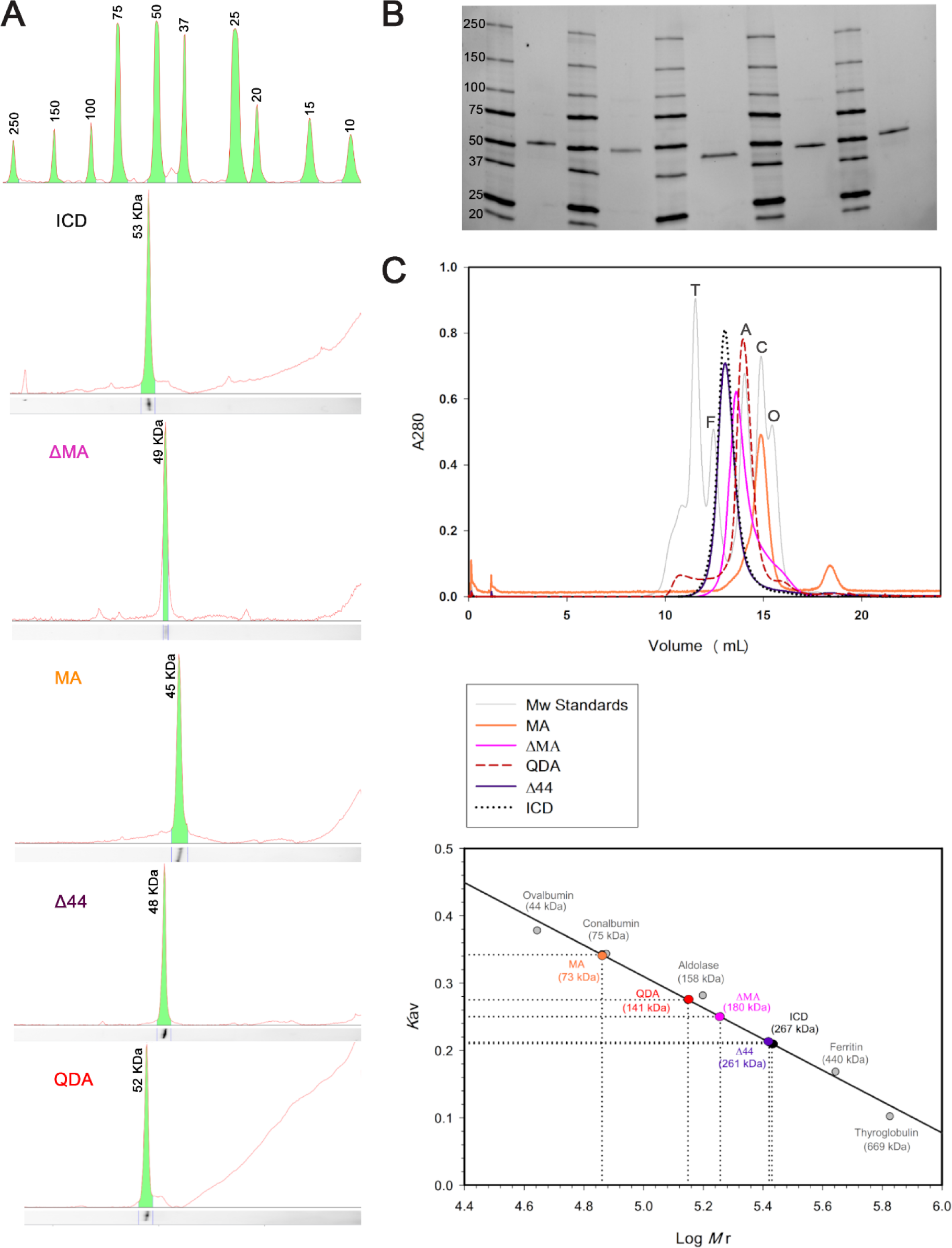
**A)** The weight of the ICD chimeras in monomeric state was determined using SDS-gel electrophoresis Molecular weight standards on top, and ICD (53 kDa), ΔMA (49 kDa), MA (45 kDa), Δ44 (48 kDa), QDA (52 kDa respectively at the bottom. **B)** The complete SDS-gel electrophoresis image (4-20% Mini-PROTEIN® TGX Stain-Free™ Biorad) from which the analysis in section A is drawn. A standard protein ladder is shown next to each protein fo increased accuracy of the analysis. Lane compositions are as followed: Lane 2: ICD, Lane 4: ΔMA, Lane 6: MA, lane 8 Δ44, and lane 10: QDA. **C)** Size-exclusion chromatography (SEC) yielded a weight of 267 kDa for ICD, 261 kDa for Δ44 180 kDa for ΔMA, 141 kDa for QDA, and 73 kDa for MA. Chromatogram of standard proteins (T, thyroglobulin; F ferritin; A, aldolase; C, conalbumin; O, ovalbumin) in grey, and chimeras in their respective colors are shown above.

### Oligomeric structure of full-length 5-HT_3A_-ICD

Affinity- or SEC-purified full-length ICD when analyzed by SDS-PAGE under reducing conditions, migrated as a major band with a relative molecular weight of 53 kDa (Figures 2 and 3) consistent with its theoretical mass of 53.4 kDa. Notably, during SEC ICD eluted much earlier than would be anticipated from its relative molecular weight determined by SDS-PAGE. The elution profile of the protein was similar over the entire range of concentrations examined (0.1 - 5 mg/ml; data not shown). The hydrodynamic volume (13.01 ml) of a well-resolved peak of ICD predicted an apparent molecular mass of 267 kDa when compared with the elution profiles of proteins with known molecular masses (Figure 3). The elution volume indicates a mass that is five times the monomer mass and therefore potential pentameric assembly. Asymmetrically shaped molecules containing multiple helical regions with an overall extended conformation, such as human mannose-binding protein, are known to display anomalous behavior on SEC (49). As shape-dependent factors can influence the elution volume of macromolecules when analyzed on SEC, the results from this hydrodynamic technique are possibly vulnerable to misinterpretation (50). Therefore, we used the shape-independent technique of multi-angle light scattering measurements coupled with SEC (SEC-MALS) to study the self-assembly properties of 5-HT_3A_-ICD constructs further (Table 1). SEC-MALS determines the absolute molecular mass of macromolecules independent of molecular shape, and the high-resolution separation by SEC facilitates analysis of each resolved protein species in a tandem manner. Examination of the SEC-MALS results revealed that wild-type 5-HT_3A_-ICD chimera exists predominantly as pentamers with an absolute molecular mass of 259 kDa (44, 45).

**Table 1:** Monomer and oligomer masses.

| Construct | Theoretical molar mass<br>of the monomer<br>($M_w$, $10^3$ g/mole) | Calculated molar mass from<br>SEC-MALS ($M_w$, $10^3$ g/mole)<br>[ Mean ± SEM ] | Dispersity ($\mathcal{D} = M_w/M_n$)<br>[ Mean ± SEM ] | Number of present monomers<br>(Calculated $M_w$ /Theoretical $M_w$) |
| --- | --- | --- | --- | --- |
| ICD | 53.4 | 259 ± 8 | 1.073 ± 0.015 | 4.8 |
| $\Delta 44$ | 49.3 | 245 ± 9 | 1.002 ± 0.001 | 4.9 |
| ΔMA | 49.8 | 149 ± 6 | 1.001 ± 0.001 | 2.9 |
| MA | 44.3 | 89 ± 2 | 1.000 ± 0.003 | 2.0 |
| QDA | 53 | 106 ± 1 | 1.004 ± 0.003 | 2.0 |

### Effect of deletions on ICD self-assembly

Our data so far demonstrate that the pentameric assembly is an intrinsic property of the ICD. An attractive possibility is that the domain-domain interaction occurs between ICDs of the neighboring 5-HT_3A_ receptor subunits. Indeed, based on the recent cryo-EM structure of the mouse 5-HT_3A_ receptor, the density map of the ICD, though partially structured, provides an inkling that the disordered stretch of the ICD between the MX and MA helices may form a ‘putative contact’ with the MA helix of the neighboring subunit (20). Moreover, an elegant mutagenesis study has implicated a cluster of three arginine residues, R432, R436 and R440 (residue numbering based on human 5-HT_3A_ subunits), within the MA helix that governs the low channel conductance of homomeric 5-HT_3A_ receptors (35). Interestingly, the same triplet arginine residues, via inter-subunit salt bridge formation with negatively charged residues in the vicinity, in particular D312 in L1, likely confer an additional tether underlying the domain-domain interaction (20, 51). Thus, by severing these inter- and/or intra- molecular interactions, we may be able to destabilize the pentameric assembly of the ICD and expect the formation of its lower-order oligomeric state(s). Therefore, we first sought to understand the effects of systematic deletions within the ICD on the oligomeric states of correspondingly derived truncated ICD chimeras.

We generated three ICD deletion constructs: 1. ICD lacking 44 amino acids from the disordered flexible region between the MX and MA helices (Δ44), 2. ICD lacking the MA helix (ΔMA) and 3. MA helix alone (MA) (Figure 1). The theoretical molecular weights, calculated from their amino acid composition, of 49.3 kDa, 49.8 kDa and 44.3 kDa, respectively were comparable to the experimentally determined relative molecular weights of 48 kDa, 49 kDa and 45 kDa, when purified protein samples were analyzed by SDS-PAGE (Figure 3).

Upon passing through a Biorad ENrich™ SEC 650 sizing column and comparing with the elution profiles of known molecular weight protein standards, the three deletion constructs showed strikingly different elution profiles (Figure 3). Importantly, all constructs eluted in single monodisperse peaks. Based on the elution volumes of 13.04 ml, 13.57 ml and 14.85 ml for Δ44, ΔMA and MA, respectively, apparent molecular weights of 261 kDa, 171 kDa, and 73 kDa were estimated. These results were inconsistent with a monomeric state for all deletion constructs. While SEC data for Δ44 may indicate a pentamer (49.3 kDa * 5 = 247 kDa vs 261 kDa), data for ΔMA and MA did not correspond to a defined multimeric state. Therefore, at this juncture, it was imperative to investigate all constructs with SEC-MALS to determine the absolute molecular weight (Table 1). The predominant peak for the Δ44 construct yielded a weight-average molar mass of 245 ± 9 kDa (mean ± S.E.M., *n* = 5), which corresponds well to a pentameric assembly (theoretical molecular mass 49.3 kDa). Moreover, the pentameric protein is monodisperse as the experimentally determined molar masses were equivalent across the peak, as indicated by the measurements of polydispersity: *M*_w_/*M*_n_ of 1.002 ± 0.001, where *M*_w_ is the weight-average molar mass and *M*_n_ is the number-average molar mass (Figure 4; Table 1). Similarly, for the other two deletion constructs, ΔMA and MA, analysis yielded the weight-average molar masses of 149 ± 6 kDa (mean ± S.E.M., *n* = 3) and 89 ± 2 kDa (mean ± S.E.M., *n* = 3), and polydispersity of 1.001 ± 0.001 (mean ± S.E.M., *n* = 3) and 1.000 ± 0.003 (mean ± S.E.M., *n* = 3), respectively. Therefore, SEC-MALS results established that ΔMA is a trimer (theoretical molecular mass 49.8 kDa) and MA is a dimer (theoretical molecular mass 44.3 kDa) in solution. It is also important to note that we did not observe a plateau in the molar mass profiles or a peak in the elution profiles that corresponded to major quantities of monomeric form of either of the deletion constructs. Together, the results suggested that we were able to disrupt pentameric assembly of the ICD. Since Δ44 assembled into pentamers, but both ΔMA and MA did not do so, we inferred at this point that the segment removed in Δ44 is not involved in pentamerization, whereas motifs contained in both MA and ΔMA may interact to mediate pentamerization.

**Figure 4.**
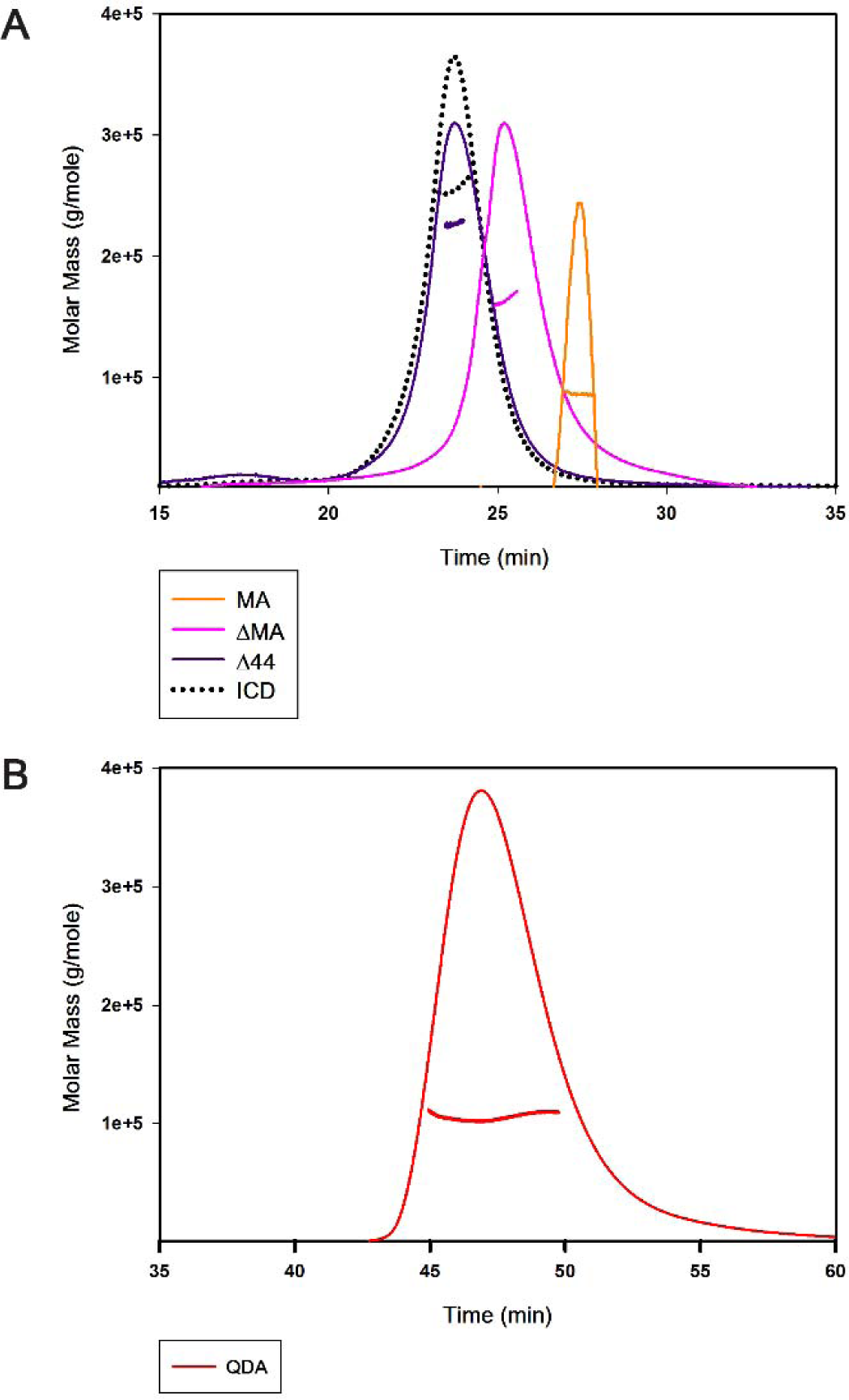
SEC-MALS determination of oligomeric assembly of ICD chimeras: **A)** The ICD with the weightaverage molar mass of 259 kDa forms a pentamer (dotted black); the Δ44 with the weight-average molar mass of 245 ± 9 kDa (purple line) retains the pentameric assembly; the ΔMA with the weight average molar mass of 149 ± 6 kDa (Pink line) and the MA with the weight average molar mass of 89 ± 2 kDa (orange line) disrupt the pentamerization of the ICD. **B)** The QDA mutation with the weight-average molar mass of 106 ± 1 kDa (red line) disrupts the pentameric assembly of the ICD. A representative SEC-MALS profile showing the Rayleigh ratio in corresponding colors for each chimera is included and the weightaverage molar mass is computed from light scattering. Note that different flow-rates were used during the monitoring period for A (0.5 ml/min) and B (0.3 ml/min).

### Effect of specific amino acid substitutions in the MA helix on the oligomeric structure of the ICD chimera

The observed involvement of both the proximal and distal segments of the ICD in pentamerization was reminiscent of the salt-bridge networks identified in some of the structural studies. Three arginine residues of the MA helix (R432, R436, R440) that have been identified as determinants for the 5-HT_3A_ subunit’s low single channel conductance participate in the configuration of putative noncovalent interactions with acidic amino acids, within 4 Å radius, from neighboring ICDs, at physiological pH, to form stabilizing salt-bridges (14, 35). We simultaneously replaced these three arginine residues, with the aligned residues glutamine, aspartic acid and alanine of 5-HT_3B_ subunits to obtain the QDA construct (R432Q/R436D/R440A) to investigate their impact on the pentamerization of the ICD.

The QDA construct eluted, at a volume of 13.92 ml, as a major symmetrical peak after SEC which predicted its apparent molecular weight as 137 kDa (Figure 3). The size of the monomer, calculated from its amino acid sequence as well as determined by SDS-PAGE, is 53 kDa (Figure 3). Analogous to the experimental strategy employed for the deletion constructs, we determined an absolute molar mass for the QDA construct by SEC-MALS. The result of the weight-average molar mass of 106 ± 1 kDa (mean ± S.E.M., *n* = 3), polydispersity of 1.004 ± 0.003 (mean ± S.E.M., *n* = 3) suggested that the QDA construct exists as a dimer in solution (Figure 4). This result indicates that mutation of the RRR motif and the likely resulting disruption of salt bridges is sufficient to abolish pentameric assembly of the ICD.

## DISCUSSION

The intracellular domain (ICD) is the most diverse and the least understood portion of pentameric ligand-gated ion channels (pLGICs) across the Cys-loop receptor superfamily. The native coding sequence of the ICD of 5-HT_3A_ receptors (5-HT_3A_-ICD), a cationic pLGIC member of the superfamily, translates into about 115 amino acids. The amino acid sequence of the 5-HT_3A_-ICD when analyzed by secondary structure prediction algorithms (PSIPRED), indicates two α-helical regions, one α-helix of 19 residues (MX) after a short linker following the M3 transmembrane helix, and a second α-helix of 33 residues (MA) continuous with and preceding M4. In the *Torpedo* nAChR structure the five MA-helices were viewed as an inverted cone protruding into the cytosol, and enclosing the intracellular vestibule of the transmembrane pore (22). Several studies show that the ICD plays a pivotal role in receptor assembly, trafficking, targeting (53, 54), functional interactions with cytoplasmic proteins (37, 55), gating and desensitization (56), inward rectification (57), Ca^2+^ permeability (58), and single-channel conductance (35, 59). In case of the 5-HT_3A_-ICD, it has become apparent that several of the experimentally determined functions are, perhaps most critically, contingent on the integrity of its secondary and/or quaternary structure. Mutagenesis studies have demonstrated that the variation in the length and amino acid composition of the ICD can influence the desensitization rate of the receptor channel (25, 57). The adjacent MA-helices form the characteristic narrow asymmetric ion permeation conduit through the lateral portals. A deviation toward a smaller size from the required minimum loop length connecting the M3 to the MA-helix has been shown to disrupt the portal architecture resulting in loss of inward rectification of macroscopic currents of 5-HT_3A_ receptors (57). Key residues within the MA-helix not only profoundly affect the permeability of Ca^2+^ but also the single channel conductance uniquely observed in experiments with homomeric 5-HT_3A_ receptors. The homomeric 5-HT_3A_R displays a very low single-channel conductance in the sub-pS range (60). This is in stark contrast to the larger currents (16-30 pS) measured from heteromeric 5-HT_3AB_ receptors as well as other Cys-loop receptors (61, 62). A seminal study attributed the unusually low conductance to a group of three arginine residues, which are found in the MA-helix of the 3A subunit but are absent in the 3B subunit (35). The X-ray crystallograph of the mouse 5-HT_3A_R revealed that some of the conductance-restricting arginine residues as well as other basic residues lining the MA-helix are within salt-bridge distance to several acidic residues on the neighboring MA-helices (14). Interestingly, it has been postulated that inter- and intra-subunit salt-bridges confer structural rigidity to the inverted pentacone formed by the five MA-helices, and that a controlled removal of these salt-bridges increases flexibility in this area (51). The same study proposed that it was widening of the lateral portals between the MA-helices, due to the increased flexibility in the absence of certain salt-bridges, and not the abolition of electrostatic repulsion that relieved the limited conductance of the 5-HT_3A_R channel. Importantly, the authors’ based their prediction on a cautious assumption that the mutations they introduced would not destabilize global inter-helical interactions that might further widen the portals (51). Taken together these findings suggest that the extensive inter-helical interactions between the MA-helices may be central to hold the intact quaternary structure of the ICD, and thus contribute to functional landscape of full-length 5-HT_3A_Rs. In this regard, it is intriguing to find that the 5-HT_3A_-ICD is structured as pentamers in the absence of both the ECD and the TMD (44). Multiple studies underscore the importance of salt-bridge formation in the stabilization of a protein oligomeric assembly (63-65). Therefore, we interrogated whether some of the aforementioned function-governing salt bridges additionally play a fundamental role in pentameric assembly.

We previously developed a chimeric strategy, where the 5-HT_3A_-ICD was linked N-terminally to a maltose-binding protein (MBP) (44, 45). In the present work, we systematically introduced deletions or defined amino acid substitutions within this construct. Firstly, in order to obtain evidence of the role of the structural change in the flexible unresolved region of the ICD, we removed 44 amino acids from this region (Δ44). A similar construct was generated in full-length channels in a study which investigated the significance of the lateral portals within MA-helices for inward rectification observed in 5-HT_3A_Rs (57). Using a series of truncation mutants, the authors determined the minimum number of residues necessary to link M3 with the MA-helix without abolishing function. While truncations of 44 and 24 amino acids from the ICD produced functional receptors, the construct with 44 amino acids deleted exhibited loss of rectification. This indicated reduced impediment to ion conduction that was inferred to be caused by deconstruction of the lateral portals. Interestingly, our results showed that truncation of the exact 44 amino acids from the soluble ICD maintained pentamers in solution. The study additionally reported a truncation of 55 amino acids in L2, or removal of 10 amino acids from the proximal part of the ICD that did not produce functional homomeric 5-HT_3A_ receptors (57). We infer that the large 55 residue truncation as well as those made in the proximal ICD can herald a global disruption and lead to the formation of a global non-native ICD conformation which may further impose unfavorable constraints for channel translocation. Our assumption is supported by the previously reported data where an engineered 5-HT_3A_R from the HEK-293 cell membrane, containing a heptapeptide replacing the entire ICD, was not only fully functional but also exhibited enhanced single channel conductance (23). Within this context, given our experimental evidence of intact pentamers formed by Δ44, we inferred that this shortened ICD plausibly underwent merely a local perturbation likely limited only to the lateral portals without disturbing the key interactions holding together the oligomeric assembly. This might have resulted in a minimally aberrant native-like ICD architecture responsible for the observed retention of function in correspondingly engineered 5-HT_3A_ receptors (57). In the structures there is a ∼54 Å gap between the end of MX and the start of the MA α-helices (M3-WILC to VRG-M4). At least 18 amino acids are needed for a linker to span this distance. Δ44 and Δ55 contain 14 and 3 amino acids in our constructs, respectively. Since (57) used the long variant with 6 additional amino acids in this segment, their Δ44 and Δ55 contain 20 and 9 amino acids. Consequently, it would be expected that Δ44 has a sufficient remaining L2 loop length to not disrupt the relative arrangement of MX and MA, and indeed it pentamerizes. Δ55, on the other hand, has a too short L2-linker and a drastic distortion is expected, leading to the disruption of pentamers (not shown). Collectively, we postulated that the underlying molecular interactions driving the ICD pentamerization must lie within the proximal (L1-MX) as well as the distal (MA) regions of the ICD.

To test the above stated hypothesis, we decided to explore the effect of deletion of a proximal (MA construct) or a distal portion (ΔMA construct) of the ICD on its oligomeric state. SEC-MALS indicated disruption of pentameric assembly in both constructs, MA was determined as a dimer and ΔMA as a trimer. The fact that the MA-construct did not pentamerize was somewhat surprising since the MA-helices are tightly associated in the crystal structure where rings of hydrophobic residues L402, L406, I409 and L413 shape a 17-Å-long narrow channel (14). Given the observation that ΔMA also disrupts the pentamerization, while Δ44 retains the pentameric state, we hypothesized that the elements of both MA and ΔMA are required for driving the pentamerization. In particular, the results pointed towards both the proximal and distal segments of the ICD as being determinants for pentameric assembly. Interestingly, when we previously made chimeras by using the prokaryotic pLGIC from *Gloeobacter violaceus* that naturally lacks an ICD and inserted ICDs from different eukaryotic anion- or cation-conducting channels, we found that the linking amino acids, both on the proximal and distal insertion sites, were crucial for functional 5-HT_3A_-ICD insertion (66). Remarkably, for other ICDs that we investigated (nAChα7, Glyα1, GABAρ1) the linking amino acids for the ICDs had little impact on function, indicating a unique characteristic for the 5-HT_3A_-ICD (67). We next sought to examine if the conductance-limiting arginines are additionally involved in maintaining the quaternary pentameric structure of the ICD. Although the arginine side chains are not well resolved in the crystal structure of the mouse 5-HT_3A_R, they can be seen enveloped by acidic residues (14), a favorable condition known to be conducive for extensive salt-bridge formation (51). Interestingly, the location of the RRR coincides with a kink in the paths of the long continuous MA-M4 helices that collectively leads to a significantly tighter packing of MA-helices as they traverse away from the membrane. The bottom aperture of the MA bundle has a diameter of 4.2 Å, too small for conduction of hydrated cations, but wider than the remaining opening of the lateral windows through which the post-M3 loop is threaded (14). Combined replacement of RRR with QDA subunit substantially augments the single channel conductance (35), a phenomenon originally attributed to the removal of repulsive forces of the positive charges of the arginines upon conducted cations. On the other hand, constrained geometric simulations, upon abolishing certain salt-bridges, have predicted enhanced structural flexibility within the MA-helices, and led to postulate that a rigid structural framework of the inverted cone formed by the MA-helices might be crucial for the limited conductance of the 5-HT_3A_ receptor channel (51). Interestingly, the QDA substitutions in our soluble construct disrupted the pentameric state of the full-length ICD and reduced it to dimers. Clearly, this finding does not appear to support the ‘charge-repulsion theory’, but rather provides the first experimental evidence of structural implications of the disjunction of salt-bridges due to the QDA substitutions with the resultant disruption of the interactions which are critical to sustain tight packing of the MA bundle as well as pentameric assembly of the ICD.

We do not know exactly why the ΔMA construct forms trimers under the given experimental conditions nor do we know the exact driving forces behind dimerization of the MA and QDA constructs in solution. It is a possibility, as documented elsewhere (65), that the breakage of pre-existing salt-bridges can induce protein remodeling via initiation of partial unfolding and reorganization of electrostatic interactions leading to the formation of non-native oligomers. Nevertheless, it is important to emphasize that we did not observe pentamers formed by any of the engineered proteins during our investigation, with the notable exception of ICD and Δ44.

Finally, to the best of our knowledge, this is the first experimental evidence which extends our understanding beyond the known functional contributions of the three arginine residues to limit conductance, and posits that their existence within the MA-helix of the 5-HT_3A_R is critical for preserving a pentameric state of the ICD.

## AUTHOR CONTRIBUTIONS

M.J. designed research. A.P., E.P., A.G.S. and M.J. conceived the wild-type and engineered constructs. A.P., E.P., A.G.S. and M.J. performed experiments and analyzed data. A.P., E.P., A.G.S. and M.J. wrote the paper.

## ABBREVIATIONS

5-HT_3A_: serotonin type 3A
BSA: bovine serum albumin
DNAse I: deoxyribonuclease I
*E. coli*: *Escherichia coli*
ECD: extracellular domain
EM: electron microscopy
GABA_A_: γ-aminobutyric acid type A
ICD: intracellular domain
IPTG: Isopropyl β-D-1-thiogalactopyranoside
MBP: maltose-binding protein
M3: third transmembrane segment
M4: fourth transmembrane segment
MA: membrane-associate α-helix
MBP-5-HT_3A_-ICD: a chimera containing maltose-binding protein and the intracellular domain of serotonin type 3A receptors
nAChRs: nicotinic acetylcholine receptors
PMSF: phenylmethylsulfonyl fluoride
pLGIC: pentameric ligand-gated ion channel
SEC: size exclusion chromatography
SEC-MALS: size exclusion chromatography in line with multi-angle light scattering
TCEP-HCl: Tris(2-carboxyethyl)phosphine hydrochloride
TMD: transmembrane domain

## ACKNOWLEDGEMENTS

Research reported in this publication was supported by the National Institute of Neurological Disorders and Stroke of the National Institutes of Health under award number <u>R01NS077114</u> (to MJ). The content is solely the responsibility of the authors and does not necessarily represent the official views of the National Institutes of Health. We thank the TTUHSC Core Facilities: some of the images and or data were generated in the Image Analysis Core Facility & Molecular Biology Core Facility supported by TTUHSC.

The authors declare no conflict of interest.

---

**Supplementary Figure 1:**
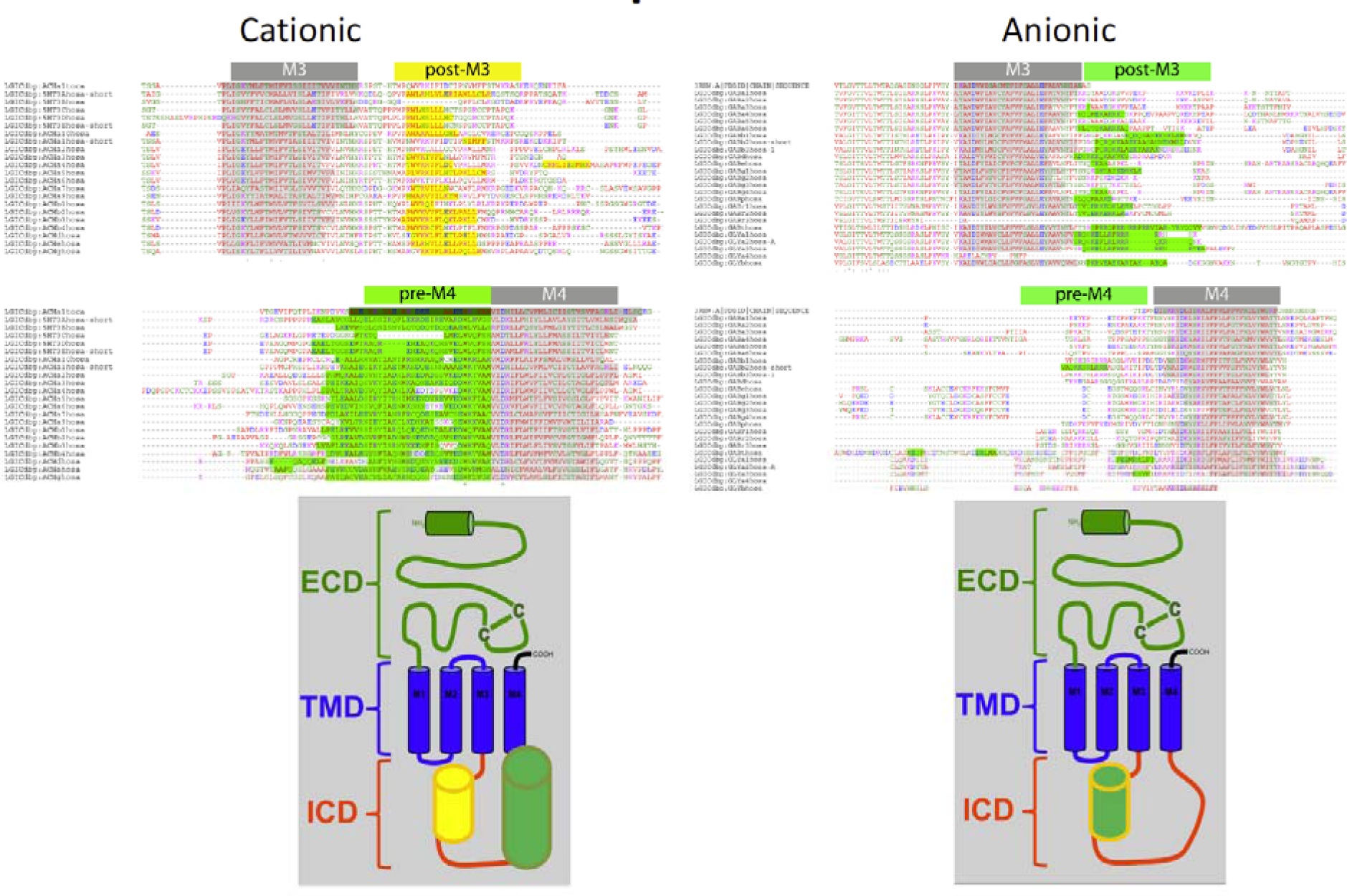
Secondary structure prediction obtained with PSIPRED. Predicted a-helical segments are highlighted in grey (transmembrane segments), yellow or green. Cation-conducting human pLGIC subunit sequences on the left and anion-conducting subunit sequences on the right.

**Supplementary Figure 2:**
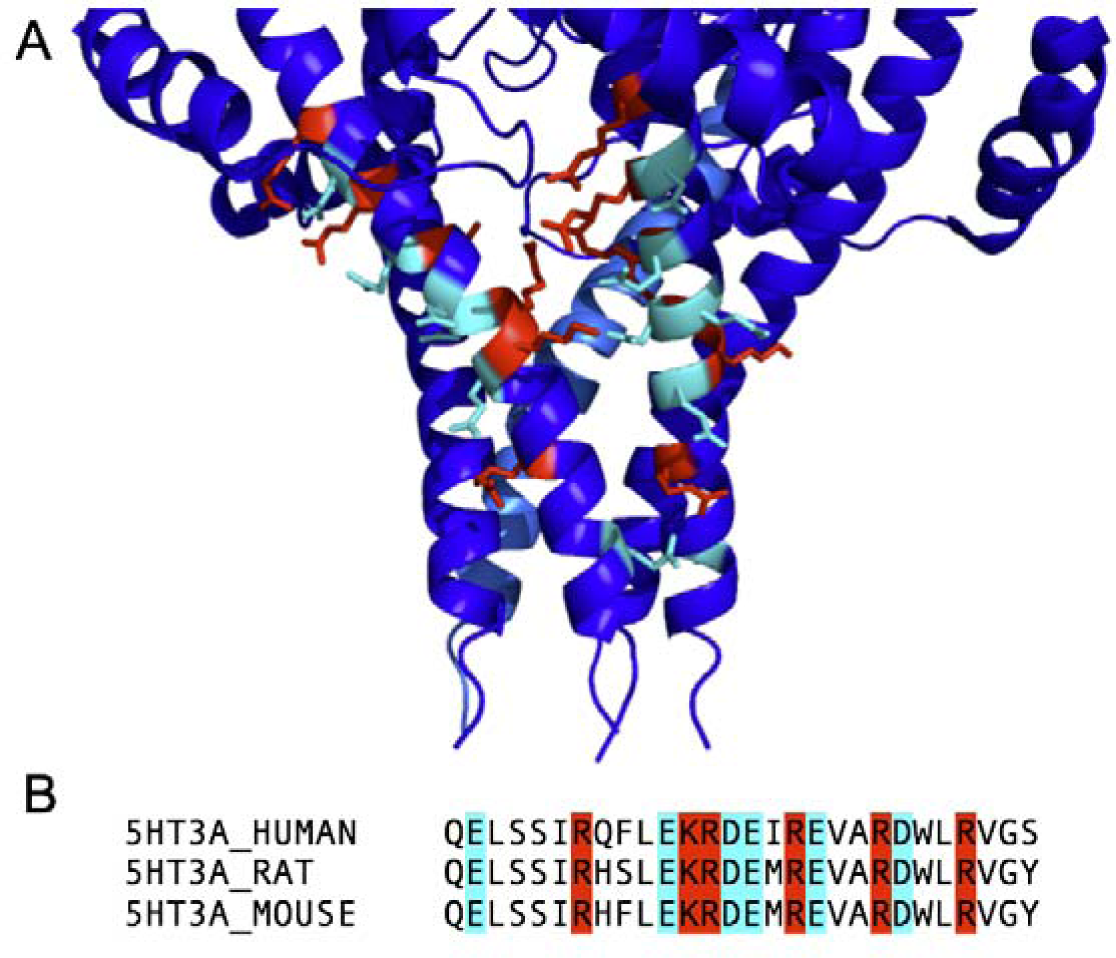
Charged amino acids in the MA-helices. A) Xray structure of m5-HT_3A_R showing amino acids potentially involved in inter-subunit salt bridges. B) Multiple sequence alignment of the MAhelix. Positive residues (red) and negative residues (cyan) are located in close proximity to each other and provide the possibility for inter-subunit salt bridges.

